# e-MutPath: Computational modelling reveals the functional landscape of genetic mutations rewiring interactome networks

**DOI:** 10.1101/2020.08.22.262386

**Authors:** Yongsheng Li, Daniel J. McGrail, Brandon Burgman, S. Stephen Yi, Nidhi Sahni

## Abstract

Understanding the functional impact of cancer somatic mutations represents a critical knowledge gap for implementing precision oncology. It has been increasingly appreciated that the ‘edgotype’ of a genomic mutation provides a fundamental link between genotype and phenotype. However, specific effects on biological signaling networks for the majority of mutations are largely unknown by experimental approaches. To resolve this challenge, we developed e-MutPath, a network-based computational method to identify candidate ‘edgetic’ mutations that perturb functional pathways. e-MutPath identifies informative paths that could be used to distinguish disease risk factors from neutral elements and to stratify disease subtypes with clinical relevance. The predicted targets are enriched in cancer vulnerability genes, known drug targets but depleted for proteins associated with side effects, demonstrating the power of network-based strategies to investigate the functional impact and perturbation profiles of genomic mutations. Together, e-MutPath represents a robust computational tool to systematically assign functions to genetic mutations, especially in the context of their specific pathway perturbation effect. The code for e-MutPath is available as a user-friendly R package at the GitHub website (https://github.com/lyshaerbin/eMutPath).

## INTRODUCTION

Genome sequencing and genome-wide association efforts have identified thousands of genetic variants across cancer types (1). Although mutations are traditionally thought to disrupt the entire gene function, it has become clear that many mutations (especially missense mutations) have a unique effect on signaling pathway perturbation (2,3), which are termed as ‘edgetic’ or ‘neomorphic’ (4–6). These mutations have been observed to exert different functional effects, including oncogenic activation and tumor suppression (6). However, the functional pathways by which genetic variants lead to diverse phenotypic consequences remain largely unresolved (7). Classical gene knockout or knockdown approaches have been performed to characterize the function of genes but not mutations (6,8,9). Indeed, it is difficult to create, express and characterize large numbers of specific mutants due to the enormous amount of time and cost involved.

Although experimental methods are critical to understand the function of individual genetic variants, it is increasingly appreciated that biological systems are formed by a large number of interacting genes (10). Interactome networks or signaling pathways exhibit distinct properties that cannot be understood by single gene-based analyses (3,11). Many mutations are believed to be edgetic, rewiring signal transduction pathways to alter cellular phenotypes (4–6). Identification of network or pathway perturbations and the consequences of such perturbations is crucial for understanding complex pleiotropic mutational effects and providing novel insights into genotype-phenotype relationships in disease (12).

While a number of methods have been developed to identify candidate driver mutations (13,14), the majority of these methods focus on the ‘nodes’ in the networks or pathways. The molecular interactions in the context of signal transduction networks are often neglected in these algorithms. To overcome this challenge, integration of available omics data from genome-scale projects is necessary, and could greatly facilitate the identification of candidate edgetic mutations and the perturbed pathways in disease (4,15). Thus, we proposed a computational method named e-MutPath (<u>e</u>dgetic <u>Mut</u>ation-mediated <u>Path</u>way perturbations), to decode the function of mutations from the pathway perturbation perspective in cancer. We applied our network-based approach to 33 cancer types and uncovered both well-known and novel cancer-associated genes and candidate driver mutations. Our results also demonstrate that e-MutPath is superior to previous methods, and more importantly, it systematically assigns functions to large numbers of genetic mutations, especially in the context of their specific pathway perturbation effect.

## MATERIAL AND METHODS

### Genome-wide genetic variants and gene expression profiles in cancer

Genome-wide somatic mutations and gene expression profile data of hepatocellular carcinoma (HCC) samples were downloaded from The Cancer Genome Atlas (TCGA) project (16). The expression of genes was measured by Fragments Per Kilobase Million (FPKM) (17), and genes not expressed in more than 30% samples were excluded in our analysis. In addition, another set of gene expression profile data for HCC was downloaded from the International Cancer Genome Consortium (ICGC) project (18). The normalized read count for each gene was used in our analysis. The gene expression profiles for another 32 types of cancer were also downloaded from the TCGA project (19) and processed in the same way as the HCC data.

### Protein-protein interaction network and signaling pathways

A systematic map of ~16,000 high-quality human binary protein-protein interaction network, which was identified by yeast two-hybrid (Y2H) assay, was used in our analysis (20,21). The homodimer interactions were removed and there are 15,957 interactions among 4,743 genes for further analysis. Human signaling pathways were obtained from (22).

### Overview of e-MutPath

The network-based method (e-MutPath) integrates genome-wide somatic mutation profiles with gene expression and a gene interaction network to identify candidate driver mutations that perturb molecular interactions (**Figure 1**). Briefly, the method includes three steps: first, perturbed gene interactions were identified in cancer based on gene expression correlation analysis; second, sample-specific interaction perturbation profiles were constructed; and finally, candidate driver mutations in each cancer sample that mediated interaction perturbations were identified by integration of interaction perturbation profiles with mutation profiles.

**Figure 1.**
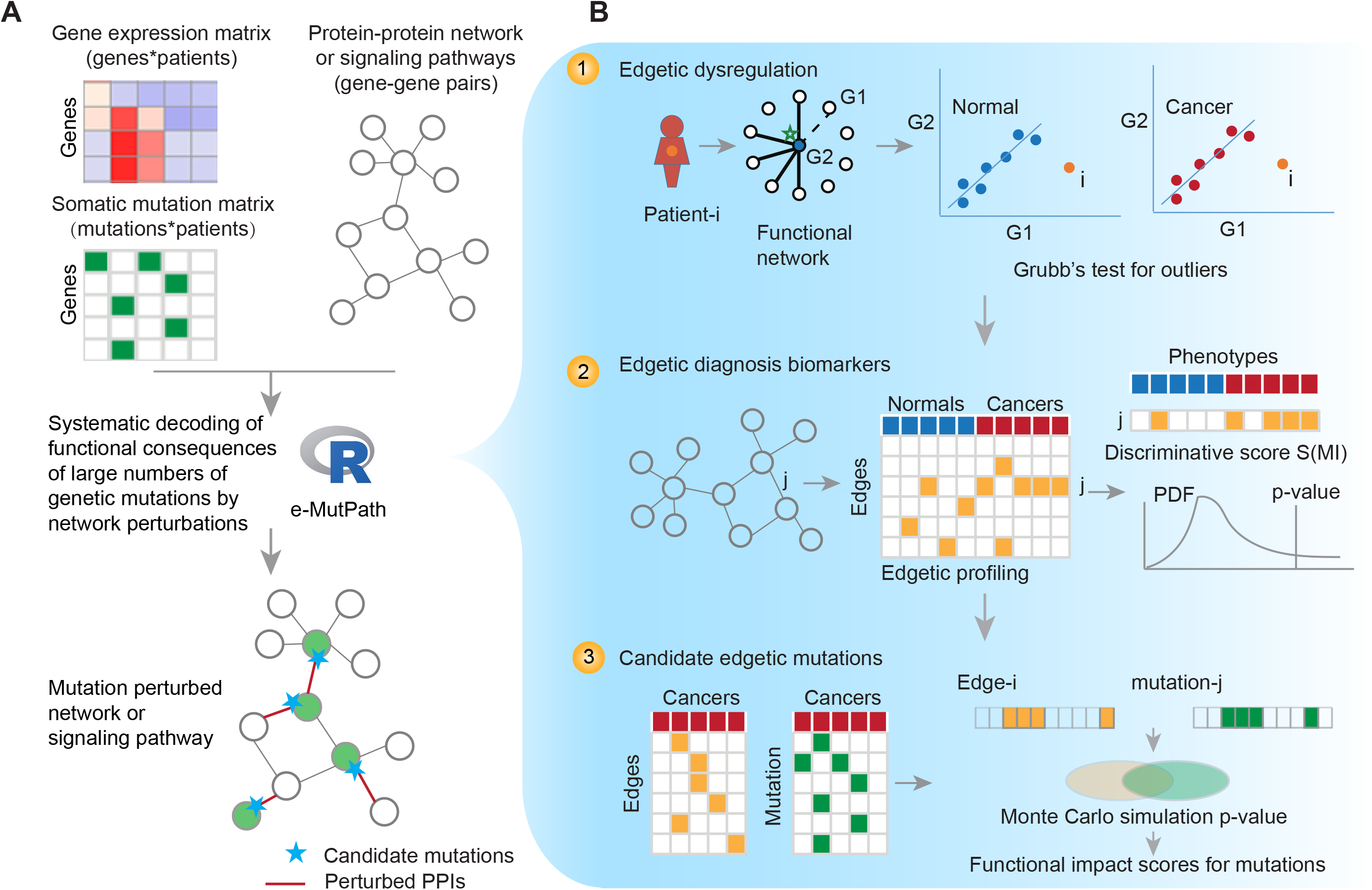
Overview of the e-MutPath. A, Gene expression and somatic mutations were integrated with functional networks to identify the candidate driver mutations. B, Identification of the mutation perturbed signaling pathway based on three steps. First, identification of patient-specific interaction perturbations based on gene expression correlation analysis. Second, identification of the edgetic biomarkers based on discriminative score. Third, identification of candidate edgetic driver mutations based on the number of overlapping patients with both mutations and interaction perturbations.

### Identifying perturbed interactions by edgetic mutations

It has been demonstrated that perturbed molecular interactions can be identified by integration of protein or gene expression profile with molecular networks (23,24). However, there are limited available protein profiles to use in complex diseases (25,26). Here, we integrated genome-wide gene expression profiles with gene interaction networks to identify sample-specific molecular interaction perturbations in cancer. The gene networks could be defined as un-directed graphG = (V, E), where V = (v_1_, v_2_, v_3_, …, v_i_, …) is the set of genes represented as nodes and E = (e_1_, e_2_, e_3_, … , e_j_, …), the set of interactions represented as edges. For each edge e_j_ formed by gene v_x_ and v_y_ in the network, we evaluated whether this edge was perturbed in sample k by calculating the distance d_k_ of this sample to other samples in the two-dimensional space of gene v_x_ and v_y_. The expression profiles of gene v_x_ and v_y_ across samples were defined as E_x_ = (e_1x_, e_2x_, e_3x_, … , e_kx_, …) and E_y_ = (e_1y_, e_2y_, e_3y_, … , e_ky_, …). The distance was calculated based on gene expression profiles as follows:

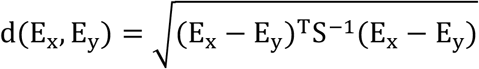

Where S is the covariance matrix of E_x_ and E_y_. When we obtained the distance of each sample, the Grubbs’ test was used to determine whether there are outliers in the distribution of the distances. Specifically, the Grubbs’ test statistic is defined as

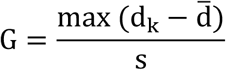

Where 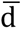 and s represent the mean and standard deviation of distance, respectively. We used the two-sided test, the hypothesis of no outliers was rejected if

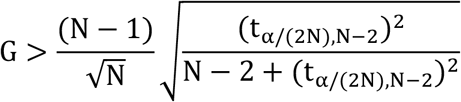

With t_α/(2N),N−2_ denoting the critical value of the t-distribution with (N-2) degrees of freedom and a significance level of α/(2N), and N is the number of samples. We detected all the outliers in the distribution, and defined these samples as exhibiting interaction perturbation in cancer. After repeating this process for all edges in the interaction networks, we identified all the perturbed molecular interactions in specific samples.

### Sample-specific molecular interaction perturbation profiles

Based on the above step, we identified all the perturbed molecular interactions in specific samples. Next, we constructed the molecular interaction perturbation profiles. Each patient was represented as a profile of binary (1, 0) states on interaction edges, where rows represented the network edges and columns corresponded to tumor samples. The elements M_ij_ in the matrix was defined as follow:

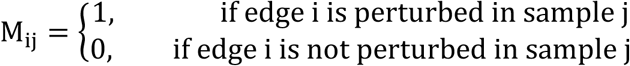

In our analysis, if there were normal samples in the gene expression profiles, we required the interaction was both perturbed when comparing with normal/cancer and among cancer samples.

### Identifying driver mutation-mediated molecular interaction perturbations

To identify candidate driver mutations that mediated molecular interaction perturbations, we firstly constructed somatic mutation profiles of all samples. The somatic mutations for each patient are represented as a binary profile, in which a ‘1’ indicates a specific mutation has occurred in the tumor. Based on the mutation profiles and molecular interaction perturbation profiles, we hypothesized that if a specific mutation and interaction perturbation co-occurred in more samples, the interaction perturbations were likely to be caused by the somatic mutation. Here, we required the mutations to occur on the genes that formed the perturbed interactions. Next, Monte Carlo simulation was used to evaluate whether a specific mutation and a perturbed interaction significantly co-occurred in tumor samples. This process was repeated 1,000 times and the p-value was defined as follows:

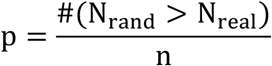

Where N_rand_ is the number of tumor samples with both the mutation and the specific perturbed interaction in random conditions, N_real_ is the number of tumor samples with both the mutation and the perturbed interaction observed, and n is the random times. Finally, we defined the functional ‘impact score’ for each mutation as 1 – p-value.

### Identification of informative molecular interactions for classification

To identify the discriminative edgetic biomarkers for classification of cancer and normal samples, we defined the discriminative score (S) for each edge in the interaction networks. Let ***a*** represent a vector of perturbation profiles for a specific edge over the tumor and normal samples, and let ***c*** represent the corresponding vector of class labels (normal or cancer). The discriminative score was defined as the mutual information MI between ***a*** and ***c***

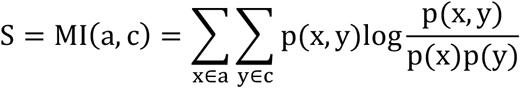

Where p(x,y) is the joint probability density function of ***a*** and ***c***, and p(x) and p(y) are the marginal probability density function of ***a*** and ***c***. x and y enumerate values of ***a*** and ***c***, respectively. Next, we tested whether the mutual information with the disease class is stronger than that obtained with random assignment of the labels to patients. This process was repeated 100 times, yielding a null distribution of MI scores. The real score of each edge is indexed on the null distribution to get the significant level. Network edges with Benjamini-Hochberg adjusted p-value<0.01 were identified as edgetic biomarkers.

### Validation of the informative molecular interaction perturbations

To validate the discriminative power of the identified edgetic biomarkers for classification of normal and cancer samples, we trained three classifiers (logistic regression, random forest and support vector machine) based on the edgetic features. To measure unbiased classification performance, we used the 10-fold cross-validation method. All the samples were divided into ten subsets of equal size. Nine subsets were used as the training set to build these classifiers using the edgetic features identified by MI score, and one subset was used as the validation set. The performance of each classifier was reported as the area under ROC curve (AUC). In addition, we constructed the classifiers based on liver cancer dataset from TCGA project and used another liver cancer dataset from ICGC as independent validation.

### Cancer subtyping analysis

Based on the interaction network perturbation profiles of liver cancer samples, consensus Non-negative matrix factorization (NMF) clustering was performed (27). To identify the subtypes, we clustered the samples by increasing k=2 to k=7 and the average silhouette width was calculated for selecting robust clusters. The clinical data for liver cancer samples were downloaded from the TCGA project and log-rank tests were performed to evaluate the survival difference among patients with different cancer subtypes. Moreover, we defined the severity score of each patient as the proportion of interactions perturbed by mutations.

### Functional analysis of informative genes

To analyze the functions of the informative genes with interaction perturbations, we used a hypergeometric test to determine whether these genes were enriched in specific pathways. The p-value was calculated as

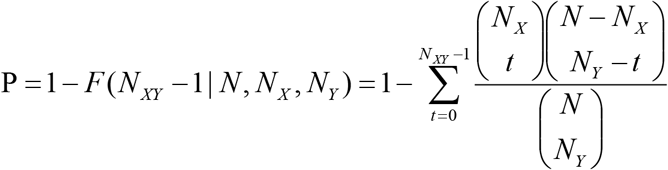

where N is the number of all genes (default background distribution), N_X_ and N_Y_ represent the total number of informative genes and genes in specific pathways, respectively, and N_XY_ is the number of overlapping genes that formed the informative edges in specific function, which had to be at least three. Kyoto Encyclopedia of Genes and Genomes (KEGG) pathways with adjusted p values <0.01 and including at least two genes were considered.

### Evaluation of the performance

To explore whether e-MutPath could retrieve cancer-related genes, we firstly obtained 602 well-known cancer-associated genes in Cancer Gene Census (28). Moreover, we downloaded the cancer-related genes from the CancerMine resource (29), a text-mined and routinely updated database of drivers, oncogenes and tumor suppressors.

The proportion of cancer-related genes, oncogenes, tumor suppressors and drivers were calculated separately for e-MutPath. In addition, we compared this proportion to four widely-used computational methods, including SIFT (30), Polyphen-2 (14), MutationTaster (13) and MutationAssessor (31). For each mutation, we calculated these four scores and the deleterious mutations predicted by these four methods were obtained. In SIFT, mutations were predicted as deleterious based on SIFT scores (<0.05). For Polyphen-2, “probably damaging” was called if the score was between 0.909 and 1. In MutationTaster, mutations predicted as “D” (“disease_causing”) were considered as driver mutations, and in MutationAssessor, mutations predicted as functional (H, M) were considered disease-relevant. All these mutations were mapped to genes and the proportion of cancer-related genes was calculated.

### Experimental validation of mutation-perturbed PPIs

We implemented a site-directed mutagenesis pipeline to generate specific mutations as in our previous studies (5,32). The mutation clones were sequenced by GENEWIZ and the sequences were confirmed based on the Basic Local Alignment Search Tool (BLAST) programs. We firstly checked whether the sequences were blasted to the exact genes of interest and then verified whether the mutations were successfully cloned into the plasmids. We next used the yeast two-hybrid (Y2H) assay to interrogate mutation-induced PPI alterations based on our previous pipeline (33). All protein interaction assays were performed twice independently.

### Data availability statement

All data presented are available and described in Online Methods. The codes for e-MutPath are freely available at the GitHub (https://github.com/lyshaerbin/eMutPath).

## RESULTS

### Overview of e-MutPath

To identify the mutations that might perturb signaling pathways, we hypothesized that gene-gene relationships would show perturbations in patients with specific driver mutations. We therefore developed e-MutPath as an open-source R package to identify candidate driver mutations that perturb functional pathways. Three types of omics datasets were integrated, including gene expression, somatic mutations and functional networks or pathways (**Figure 1A**). The output would provide prioritized mutations as well as the perturbed edges in signaling pathways or networks.

Specifically, three steps were performed in this computational method in the context of cancer. First, perturbed functional interactions were identified in each cancer patient based on a correlation perturbation analysis of RNA expression (**Figure 1B**, top panel). All the patients were mapped to a two-dimensional plane based on the expression levels of two interacting genes. If a patient showed a significant deviation from a normal gene-gene relationship distribution, the patient would be an outlier in the regression line modeled by all the patients. We used Grubb’s test for detecting the outliers (see “Methods”). Second, sample-specific interaction perturbation profiles were constructed; if gene expression data in normal samples were present, we also identified the perturbed functional subnetworks or pathways that could distinguish cancer from normal samples (**Figure 1B**, middle panel). Finally, candidate driver mutations in each cancer sample that mediated interaction perturbations were identified by integration of interaction perturbation patterns with mutational profiles (**Figure 1B**, bottom panel). We used Monte Carlo simulation to evaluate whether the patients with specific mutations were significantly overlapped with those showing gene-gene relationship perturbations (see “Methods”). The mutations with p-value less than 0.05 were identified as candidate driver edgetic mutations.

### e-MutPath identifies informative paths to distinguish disease

As a proof of principle, we applied the proposed method to hepatocellular carcinoma (HCC) data obtained from The Cancer Genome Atlas (TCGA) project (16), which included 374 cancer and 50 normal samples. Based on the human protein-protein interaction (PPI) networks, we identified 477 interactions among 486 genes perturbed in HCC (**Figure S1A**). We found that the majority of these interactions involved genes that had been demonstrated to play critical roles in cancer, such as TRIP13, MDFI, SPRY2 and MEOX2. Particularly, TRIP13 was found to promote cell proliferation, invasion and migration in cancer (34). In addition, MEOX2 was identified as a target gene of the TGF-beta/Smad pathway and was known to regulate cell proliferation (35). To systematically understand the function of the perturbed PPIs, we performed functional enrichment analysis based on the genes involved. Functional enrichment analysis revealed that the perturbed functional subnetworks were enriched in HCC-related functions, such as insulin signaling pathway (36) (P=2.29E-5) and FoxO signaling pathway(37) (P=0.001, **Figure S1B**).

Next, we investigated whether the perturbed PPI profiling could provide information for cancer subtype classification. We found that based on sample-specific functional perturbation profiles, cancer and normal samples were effectively distinguished from each other (**Figure S1C**). We next trained three types of machine learning methods (Support Vector Machine, SVM; Logistic regression, logR; and Random forest, RF) based on the perturbation profiling of patients and normal controls. Based on ten-fold cross-validation, we found that these classifiers exhibited the AUC (Area under the receiver operating characteristic curve) greater than 0.9 (**Figure 2A**, AUC=0.937-0.997). Moreover, these perturbed functional profiles were validated in another independent HCC dataset obtained from International Cancer Genome Consortium (**Figure 2A**, AUC=0.814-0.942). These results suggest that e-MutPath can identify the informative functional paths to distinguish disease from normal controls, which is helpful for cancer diagnosis.

**Figure 2.**
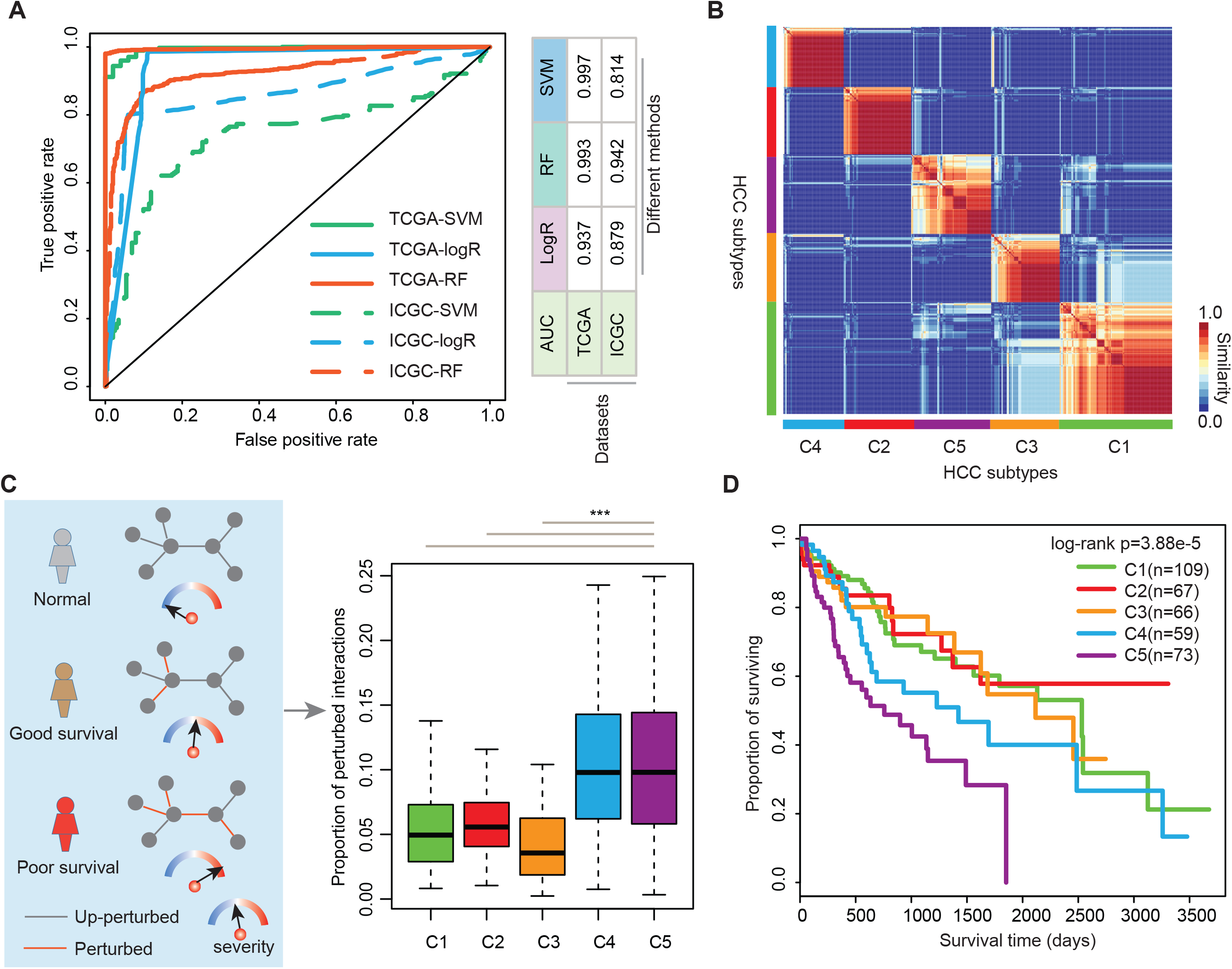
Identification of the informative signaling path based on e-MutPath in cancer. A, The ROC curve of edgetic biomarkers for classifying normal and cancer patients. The solid lines are for TCGA dataset and dash lines for ICGC dataset. Lines with different color are for three classifying algorithms. B, The heat map shows the similarity of HCC patients. Five subtypes were identified based on consensus clustering. C, The proportion of perturbed PPIs in five subtypes. D, Kaplan–Meier curve for overall survival of subtypes of HCC.

### e-MutPath stratifies cancer subtypes with clinical relevance

After identifying the perturbed PPIs in cancer, we found that some patients were clustered together (**Figure S1C**), suggesting that there were several subtypes in HCC. We thus used the interaction perturbing profiles for in-depth characterization of HCC subtypes. We stratified the HCC patients based on the similarity of their perturbation profiles by consensus clustering. Based on the k-values from 2 to 7, we identified an optimum number of five HCC subtypes (**Figure 2B** and **Figure S2A**), each consisting of 109, 67, 66, 59 and 73 patients. These subtypes were henceforth termed C1-C5. We found that there were no stage differences among these subtypes (**Figure S2B**, p=0.06).

Furthermore, we calculated the proportion of perturbed PPIs for each patient and defined this as a severity score (**Figure 2C**). We hypothesized that if the patients had higher severity scores, they would exhibit a higher risk of cancer. We found that the patients in C4 and C5 showed significantly higher severity scores (**Figure 2C**, P<0.001). To evaluate if a high proportion of perturbed PPIs could reflect disease severity in patients harboring specific mutations, we next examined the survival rates of patients. We found indeed it was the case. Patients with mutations that perturbed a higher proportion of interactions tended to correlate with a significantly shorter survival time, indicating the deleterious nature of these mutations (**Figure 2D**, log-rank P=3.88E-5). In addition, we analyzed cancer risks in patients by assessing their response to immunotherapy, which represents an alternative treatment approach that has been successful in many different cancer types (38). We found that the patients in C3 had higher immune scores, MHC scores and CYT scores than other subtypes (**Figure S2C**). These results suggest that these patients are likely to response for immunotherapy. Together, our approach is able to stratify cancer patients with distinct clinical outcomes.

### e-MutPath outperforms other methods in prioritizing cancer genes

Having shown in previous sections e-MutPath could identify mutation-perturbed signaling pathways, we next evaluated its performance in uncovering cancer-related genes in 33 cancer types. We first considered the genes from Cancer Gene Census (CGC) (39) and found that our top predictions included a high fraction of CGC genes (**Figure 3A**). To illustrate the power of e-MutPath, we compared its performance with four widely used approaches-SIFT (30), Polyphen-2 (14), MutationTaster (13) and MutationAssessor (31). For the majority of cancer types, e-MutPath consistently prioritized a larger fraction of CGC genes than other methods (**Figure 3A** and **Table S1**), demonstrating the advantage of network integration. We found that while e-MutPath predicted a smaller number of targets, they comprised larger fractions of ‘gold standard’ cancer genes by CGC (**Figure 3B**).

**Figure 3.**
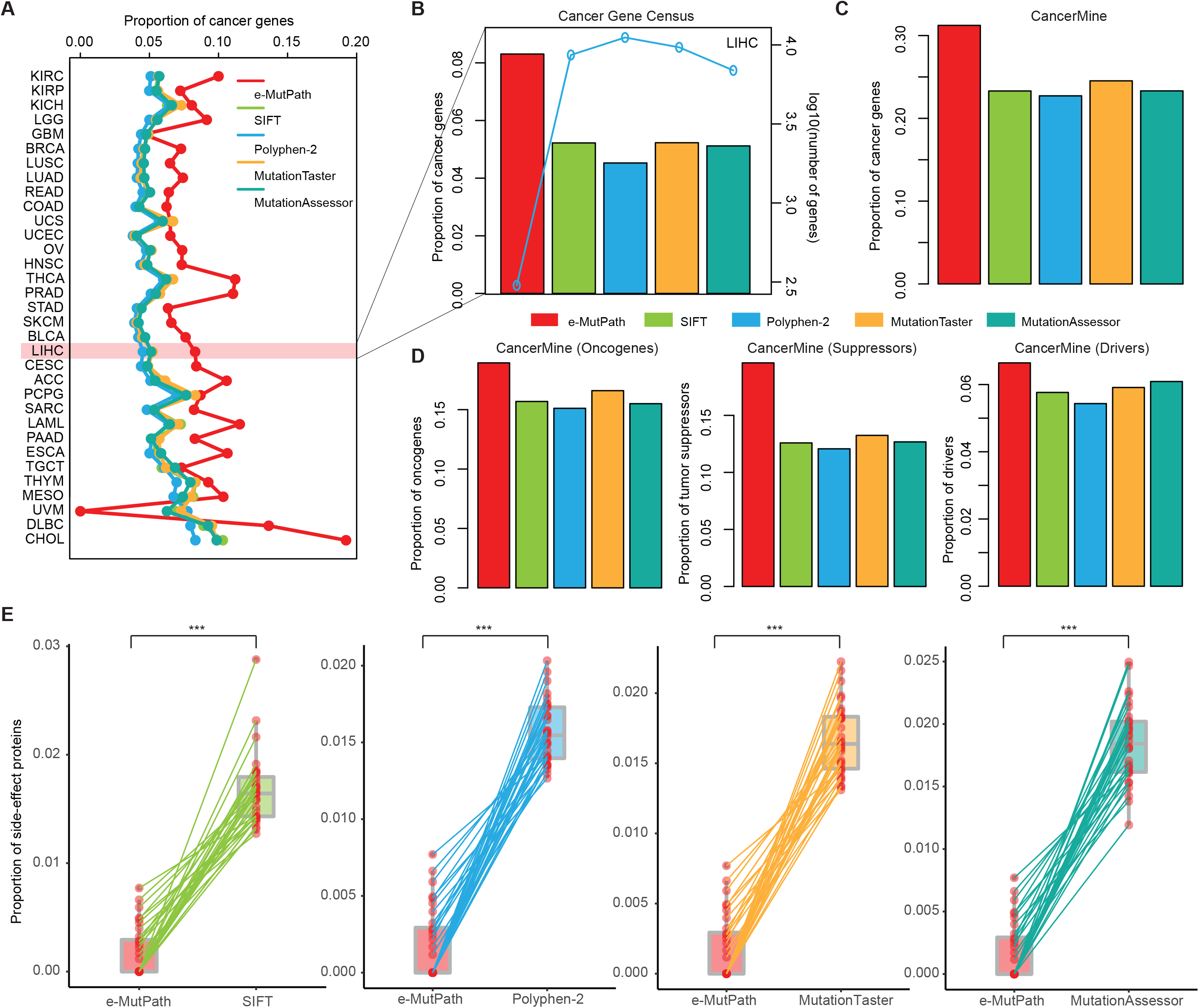
Performance evaluation of e-MutPath. A, The proportion of cancer genes identified by different methods across 33 cancer types. B, The left y-axis shows the proportion of cancer genes identified by five methods and the right y-axis shows the number of genes identified by each method. C, The proportion of cancer genes in CancerMine identified in HCC by different methods. D, The proportion of oncogene, tumor suppressors and driver genes in CancerMine identified in HCC by different methods. E, The proportion of proteins associated with side-effect identified by different methods.

In addition, we obtained known cancer genes from CancerMine (29), which is a text-mined and routinely updated database of drivers, oncogenes and tumor suppressors. We found that the overall results were consistent for all genes (**Figure 3C** and **Figure S3A**), oncogenes, tumor suppressors and driver genes (**Figure 3D** and **Figure S3B-D**). Recently, a number of large-scale screens for cancer vulnerability genes using CRISPR-Cas9 and RNAi systems have been conducted (40,41). Using these datasets, we found that the predictions by e-MutPath were more enriched for essential genes than other methods in the majority of cancer types (**Figure S4A**). Particularly, the predicted genes by e-MutPath in LUSC, BLCA and LIHC showed significantly lower essentiality scores than predictions by other methods (**Figure S4B-G**).

Finally, to evaluate potential targetable and side effects of the predicted genes, we used a list of 151 clinically actionable genes (42) and 237 proteins that are reported to be associated with side effects (43). We found that e-MutPath predictions exhibited similar fractions of actionable genes (**Figure S5**) but were depleted of side effect-causing proteins across cancer types (**Figure 3E**). Taken together, these results demonstrate significant improvement of e-MutPath over previous state-of-the-art methods in identifying cancer-related genes.

### Application of e-MutPath in pan-cancer

In order to better understand mutation-perturbed signaling pathways in cancer, we applied e-MutPath to 33 cancer types. We identified a connected network perturbed by mutations (**Figure 4A** and **Table S2**), including a number of genes (such as MED4, TCF4, and EWSR1) known to be involved in cancer. Cancer genes often function as network hub proteins which are involved in many cellular processes (44). We next investigated the topological features of the prioritized genes with mutations and found that they showed significantly higher degrees, betweenness and closeness (**Figure 4A**, p-values<0.001). Moreover, we performed functional enrichment analysis and found that these genes were likely to be involved in cancer hallmark pathways (**Figure 4B**). Collectively, these results suggest that these mutations play critical roles in cancer by perturbing the interactions involved in cancer-related pathways.

**Figure 4.**
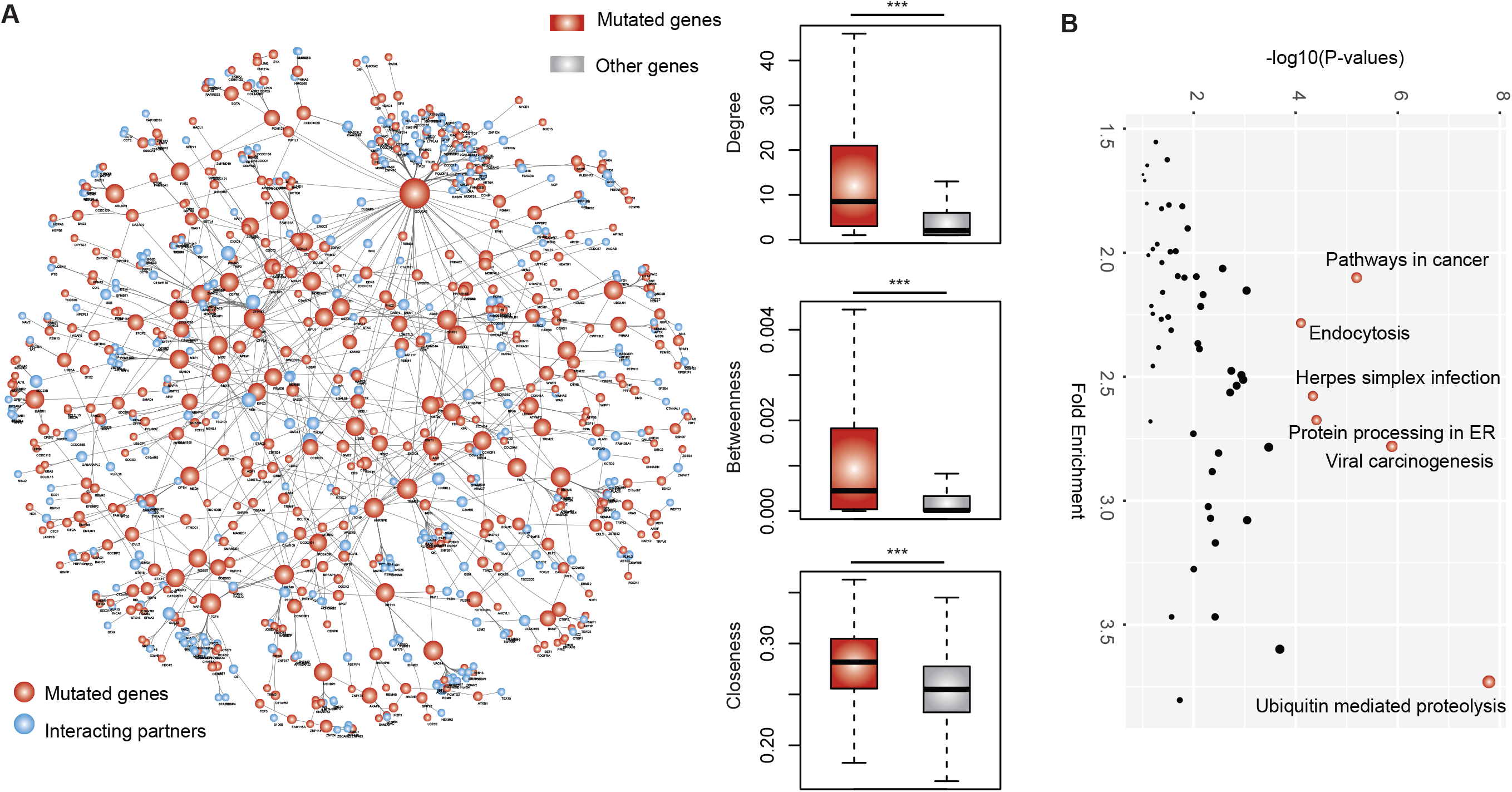
Interaction perturbations across cancer types. A, The network shows the perturbed interactions across 33 cancer types by mutations. Red, mutated genes; blue, interacting partners. The box plots show the topological features distribution for mutated genes and other genes in the original PPI network. Top, degree; middle, betweeness; bottom, closeness. B, The enriched pathways by the mutated genes.

Particularly, we found that a candidate gene KEAP1 and the interacting partner DPP3 were highly expressed in HCC compared to normal tissues (**Figure 5A**). Moreover, we investigated the correlation of gene expression with patients’ survival time. We found that high expression levels of the interacting partner gene were slightly associated with poor prognosis in HCC (**Figure 5B**). However, we found that the combined KEAP1-DPP3 signature could significantly distinguish the patients with different survival (**Figure 5B**, P=0.02). e-MutPath identified one missense mutation in KEAP1 (G379V) that perturbed the interaction between KEAP1 and DPP3 in HCC. The prediction was further validated by yeast two-hybrid assay. Lines of evidence have implied that the p62-Keap1-Nrf2 axis plays an important role in tumorigenesis (45,46). Collectively, our results suggest that these mutations play critical roles in cancer by perturbing the interactions involved in cancer-related pathways.

**Figure 5.**
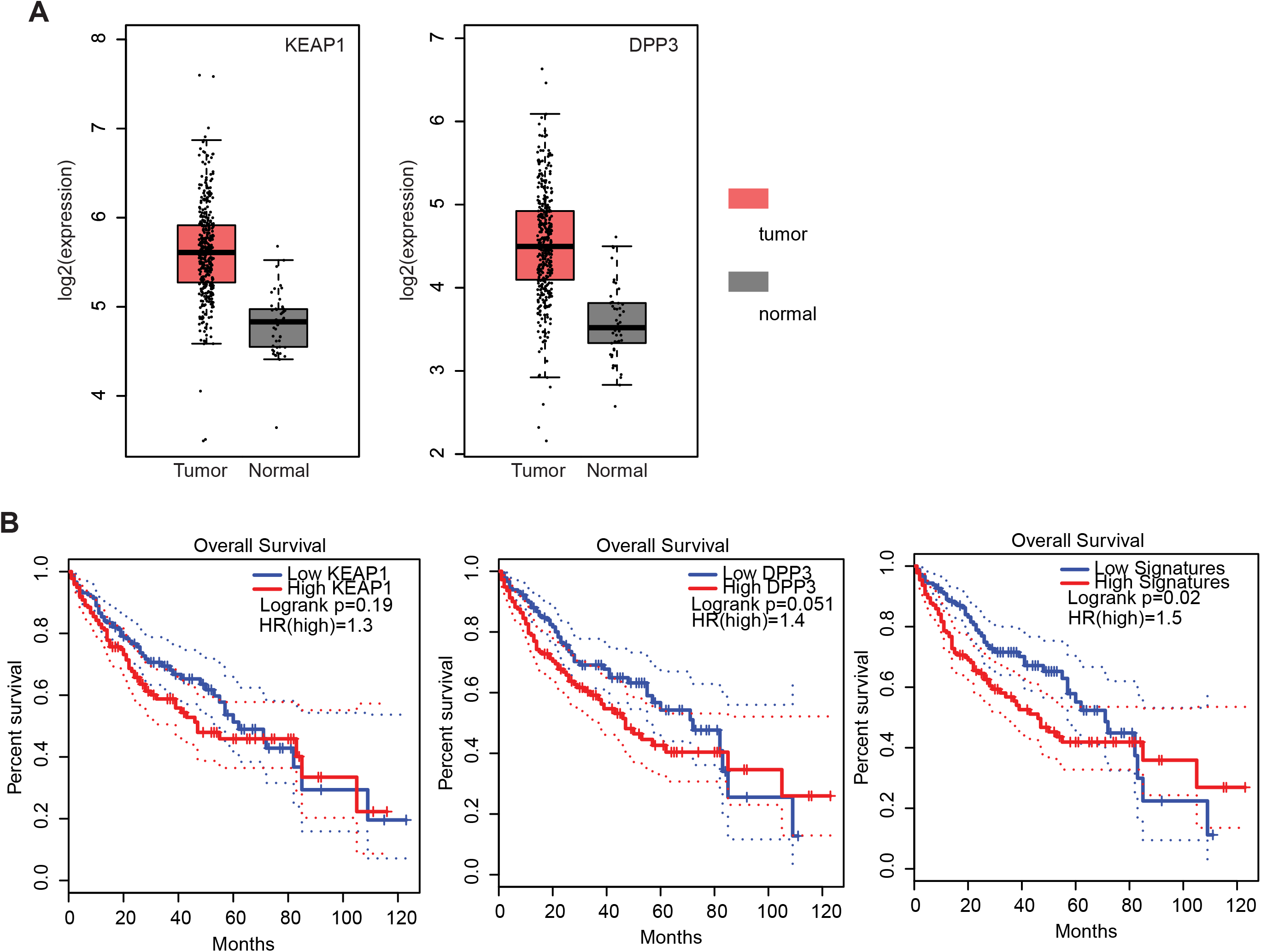
Perturbation of interactions between KEAP1 and DPP3 by mutations in HCC. A, The distribution of KEAP1 and DPP3 expression in cancer and normal samples. Red, tumor; gray, normal. B, Kaplan–Meier curve for overall survival for patients with KEAP1, DPP3 expression and combination.

### Signaling pathway perturbation in cancer

To further assess if the perturbed pathways by edgetic mutations were functionally relevant in specific disease context, we next integrated mutational profiles, gene expression patterns and signaling pathways to identify the perturbed pathways in HCC. Based on e-MutPath, we identified the top ranked HCC mutations which were mostly enriched in centrosomal genes (CEP70, CEP76 and CEP135), JAK1, EGFR and HNF1A (**Figure 6A**). JAK1 encodes a cytoplasmic tyrosine kinase that is associated with a variety of cytokine receptors and plays a critical role in cell proliferation, survival, and differentiation (47). The mutated forms of JAK1 might alter the activation of JAK/STAT pathways, and thus contribute to cancer development and progression. Moreover, we identified the hotspot in-frame mutation EGFR A767_V769dup, which perturbed the EGFR receptor tyrosine kinase signaling pathway in HCC (**Figure 6B**). The EGFR signaling pathway had been shown to play a key role in chronic liver damage, as well as in cirrhosis and HCC (48). Our results highlighted the importance of EGFR mutations in the development of liver diseases. Moreover, we identified five mutations in HNF1A ranked in the top 50 hits, which perturbed the protein interactions among HNF1A, MAPK8, PPARD and IGFBP1 (**Figure 6B**). The dysregulation of the HNF1A and PPARD signaling had been demonstrated to play fundamental roles in HCC (49). In summary, e-MutPath not only identifies critical mutations in cancer, but also provides a systems-level mechanistic view to illustrate the functions of large numbers of candidate driver mutations.

**Figure 6.**
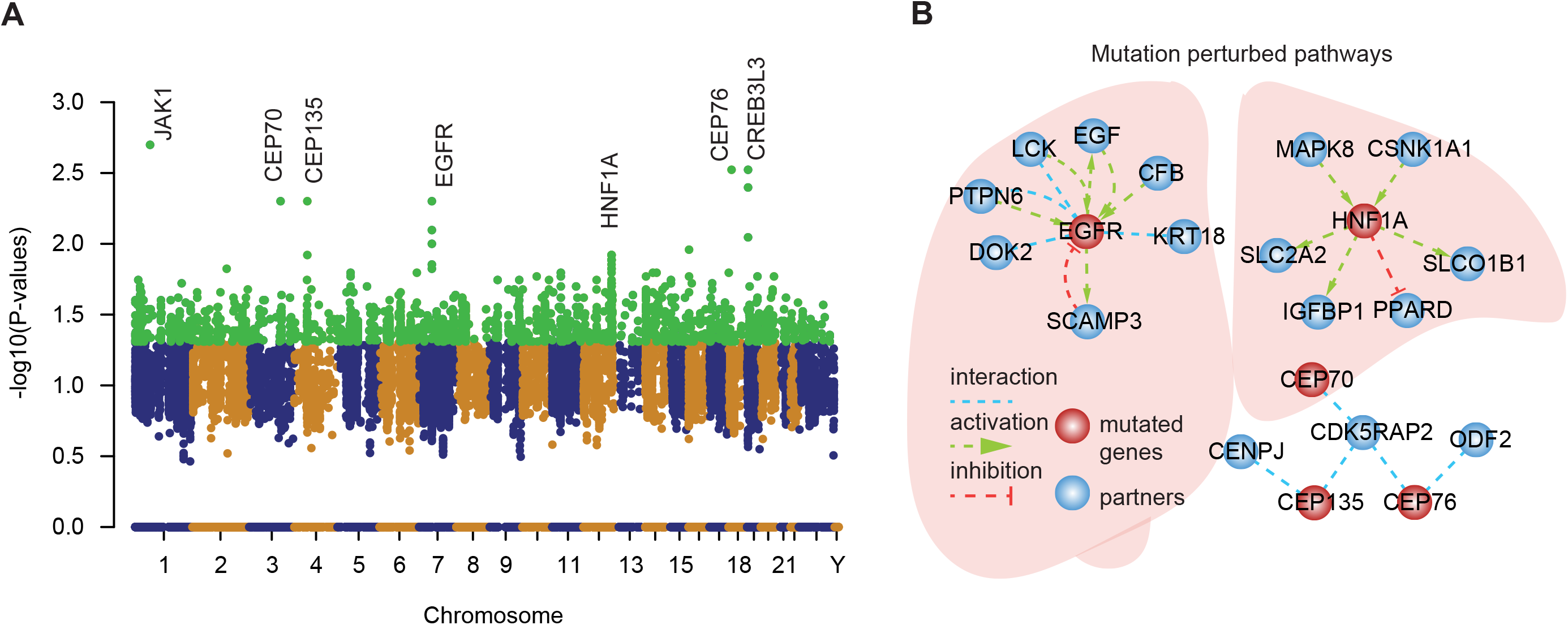
e-MutPath identified the perturbed signaling pathway mediated by mutations. A, Manhattan plot of the mutations in HCC. B, Top ranked mutations perturbed signaling pathway in HCC.

## DISCUSSION

In this study, we introduced a network-based method, e-MutPath, for decoding the functional consequences of mutations by network or pathway perturbations. This method incorporates individual mutational profiles with interaction networks or signaling pathways, which is a powerful approach for uncovering cancer-related genes. Our method is based on the ‘edgotype’ concept proposed recently and therefore complements existing frequency-based or node (protein)-based methods (50,51). We have demonstrated by the wide application of e-MutPath to 33 cancer types that, functional mutations disrupt interactions involving genes previously implicated in the development and progression of cancer, providing complementary evidence for their functional impact. The identified genes are also depleted in proteins that are associated with side-effect. Even more interesting, we found that e-MutPath could identify the perturbed pathways to stratify patients into different cancer subtypes. These results suggest that e-MutPath is a versatile tool that provides an effective framework for functional characterization of mutations in cancer and with immediate clinical relevance.

In the future, e-MutPath can be extended in a number of ways. For example, while it currently analyzes only mutations within genes, other alterations are also observed in cancer. Alterative splicing and gene-fusions have been found to perturb signaling pathways (52,53), considering both types of alterations will increase the power of our approach. Second, e-MutPath may also benefit from incorporating the development of sample-specific network. While we currently consider the interaction networks or signaling pathways, considering the context specificity of these networks or pathways may be derived from recently published methods (54). Moreover, we have demonstrated e-MutPath to identify the perturbed PPIs and signaling pathways. It can also extend to other regulatory networks, such as RNA binding protein regulatory networks, miRNA-gene regulatory networks (11,55). We have applied e-MutPath across 33 cancer types and have shown that it is broadly effective in identifying cancer-related genes. However, cancers of the same tissue can often be grouped into distinct subtypes based on molecular features (16). With the development of high throughput sequencing technology or single cell sequencing (56), e-MutPath could be used to study how different subtypes of a type of cancer yield different network or pathway perturbations and decode the function of mutations in the subtype context.

Collectively, e-MutPath could systematically uncover and stratify the functional consequences of myriad genetic mutations identified in distinct patients stricken by a variety of human diseases. Our new method identifies specific candidate driver mutations as well as their perturbed functional networks or pathways by integrating functional omics datasets at the systems level. We anticipate the proposed computational method will facilitate improved biological understanding of the function of disease variants towards personalized precision medicine.

## ACKNOWLEDGEMENTS

We were also grateful to contributions from TCGA Research Network Analysis Working Group. We also acknowledge the High-Performance Computing Facility at MD Anderson, Biomedical Research Computing Facility at UT Austin and Texas Advanced Computing Center (TACC) for computing assistance.

